# A new murine *Rpl5* (*uL18*) mutation provides a unique model of variably penetrant Diamond Blackfan Anemia

**DOI:** 10.1101/2021.02.27.430069

**Authors:** Lei Yu, Philippe Lemay, Alexander Ludlow, Marie-Claude Guyot, Morgan Jones, Fatma F. Mohamed, Ghazi-Abdullah Saroya, Christopher Panaretos, Emily Schneider, Yu Wang, Greggory Myers, Rami Khoriaty, Qing Li, Renny Franceschi, James Douglas Engel, Vesa Kaartinen, Thomas L. Rothstein, Monica J. Justice, Zoha Kibar, Sharon A. Singh

**Author notes:** These authors contributed equally (Lei Yu, Philippe Lemay, Alexander Ludlow). These authors contributed equally (Zoha Kibar, Sharon A. Singh). **Corresponding author**: Sharon A. Singh, University of Michigan, 1500 E. Medical Center Drive, Ann Arbor, MI, USA.

## Abstract

Ribosome dysfunction is implicated in multiple abnormal developmental and disease states in humans. Heterozygous germline mutations in genes encoding ribosomal proteins (RPs) are found in the majority of individuals with Diamond Blackfan anemia (DBA) while somatic mutations have been implicated in a variety of cancers and other disorders. Ribosomal protein-deficient animal models show variable phenotypes and penetrance, similar to human DBA patients. The spontaneous anemia remission observed in some DBA patients occurs via unknown mechanism(s) and has not been previously described in animal models. Here we characterized a novel ENU mouse mutant (*Skax23*^*m1Jus*^) with growth and skeletal defects, cardiac malformations and increased mortality. Following genetic mapping and whole exome sequencing, we identified an intronic *Rpl5* mutation, which segregated with all affected mice. This mutation was associated with decreased ribosome generation, consistent with *Rpl5* haploinsufficiency. *Rpl5*^*Skax23-Jus*^ mutant animals had a profound delay in erythroid maturation and increased mortality at embryonic day E12.5, which improved by E14.5. Surviving mutant animals had a macrocytic anemia at birth as well as evidence of ventricular septal defect (VSD). Surviving adult and aged mice exhibited no hematopoietic defect or VSD. We propose that this novel *Rpl5*^*Skax23-Jus*^ mutant mouse will be useful to study the factors influencing the variable penetrance and anemia remission that are observed in DBA.

## Introduction

The ribosomopathies, disorders that result from disruptions in various structural and functional elements of the ribosome, have a wide range of clinical manifestations^1–3^. Among the ribosomopathies, Diamond Blackfan anemia (DBA) is the disorder most often linked to anemia. However, many nonhematopoietic phenotypes, particularly in the skeleton and heart, are also found in DBA, as is an increased risk of developing malignancies at young age^4–8^. Inherited heterozygous mutations in genes encoding ribosomal proteins (RPs) lead to the majority of DBA cases^7^. Several mechanisms for the development of anemia in DBA have been proposed including nucleolar stress, heme toxicity, reduced translation of key erythroid differentiation proteins (e.g. GATA1) and aberrant innate immune activation^9–15^.

DBA classically presents as a severe macrocytic anemia prior to the first year of life, often requiring red cell transfusions^16^. The penetrance is quite variable, with some patients developing failure of erythropoiesis prior to birth (i.e. hydrops fetalis), while others exhibit no phenotypes even into adulthood^17^. Some individuals can cycle through periods of anemia remission and relapse for unknown reasons. Remission has been defined by the DBA registry as “an adequate hemoglobin level without any treatment, lasting 6 months, independent of prior therapy”^16^. There is no known genetic modifier, sex or treatment difference between patients who attain remission and those who remain symptomatic^18^. Approximately 20% of patients enter remission by age 25, but relapses do occur after viral illnesses or pregnancy and are more common for patients with *RPL15* (*eL15*) variants for unknown reasons^19,20^. This dynamic anemia presentation has never been adequately represented in an animal model and therefore the etiology for this variable presentation and spontaneous remission from anemia remains obscure.

The existing DBA animal models include several murine and zebrafish mutants that variably mimic some DBA disease phenotypes^21–23^. *RPS19* (*eS19*), the first identified mutated gene in DBA, is also the most common, accounting for about 25% of DBA patients^7,24^. Animal models of *Rps19* haploinsufficiency exhibit a wide variety of manifestations, ranging from no phenotype to anemia, growth retardation and variable decreases in other hematopoietic lineages (platelets and white cells)^25–27^. Here, we focus on *RPL5* (*uL18*), which is the second most commonly mutated RP gene in DBA. Patients with *RPL5* mutations often have more severe presentations with increased presence of developmental defects and a lower chance of attaining spontaneous remission^7,28-31^. Somatic mutations in *RPL5* have also been linked to various malignancies^32^. Two previous studies, one describing *Rpl5* heterozygous mice and another describing *Rpl5* depletion by shRNA, were reported^33,34^. *Rpl5* haploinsufficient mice exhibited soft tissue sarcoma but no erythroid defect, while mice with *Rpl5* depletion by shRNA demonstrated mild anemia. Zebrafish with *Rpl5* knockdown using an anti-sense morpholino oligonucleotide have also been reported exhibiting various developmental and hematopoietic defects^35^.

In this report, we generate and describe what to our best knowledge is the first murine *Rpl5* mutant mouse model of DBA that demonstrates variably penetrant anemia. We propose that these mice mimic the spontaneous remission phenotype seen in the human disease and will be instructive in future studies to better understand this elusive phenomenon.

## Materials and Methods

### Generation of the *Skax23*^*m1Jus*^ mouse mutant

The *Skax23*^*m1Jus*^ mouse mutant (MGI: 3046775) with a kinked tail and small size was generated and identified as part of a chromosome 11 balancer mutagenesis screen at the Mouse Mutagenesis and Phenotyping Center for Developmental Defects at Baylor College of Medicine in Houston, Texas. The study design of this mutagenesis screen has been described in detail elsewhere^36^. Briefly, mice heterozygous for the balancer chromosome 11 were crossed with ENU-treated C57BL/6J mice. To generate individual lines of mice that inherit the same mutation, G1 animals were crossed to generated G3 offspring, which were analyzed for linkage to the chromosome 11 balancer. *Skax23*^*m1Jus*^ was identified as a dominant mouse mutant with a kinked tail and a small size that failed to segregate to chromosome 11. *Skax23*^*m1Jus*^ G3 mutants were outcrossed onto a 129S6/SvEvTac background and the resulting N1 progeny were archived as frozen sperm and subsequently recovered at The Jackson Laboratory (https://www.jax.org/jax-mice-and-services) by *in vitro* fertilization of C57BL/6J oocytes. The *Skax23*^*m1Jus*^ mutant stock was generated and maintained by matings of presumed heterozygous mice that displayed the phenotype to 129S6/SvEv mice. For genetic mapping studies, *Skax23*^*m1Jus/*^*+* mutants with a kinked tail and small size were backcrossed to 129S6/SvEv mice for five consecutive generations. Mice were examined macroscopically for the presence of a severely ‘kinked’ tail and small size (Figure 1A–B). The protocols relevant to support this study were approved by the Institutional Committee for Animal Care in Research at Baylor College of Medicine, the Research Center of Sainte Justine Hospital, Western Michigan University School of Medicine and at the University of Michigan.

**Figure 1:**
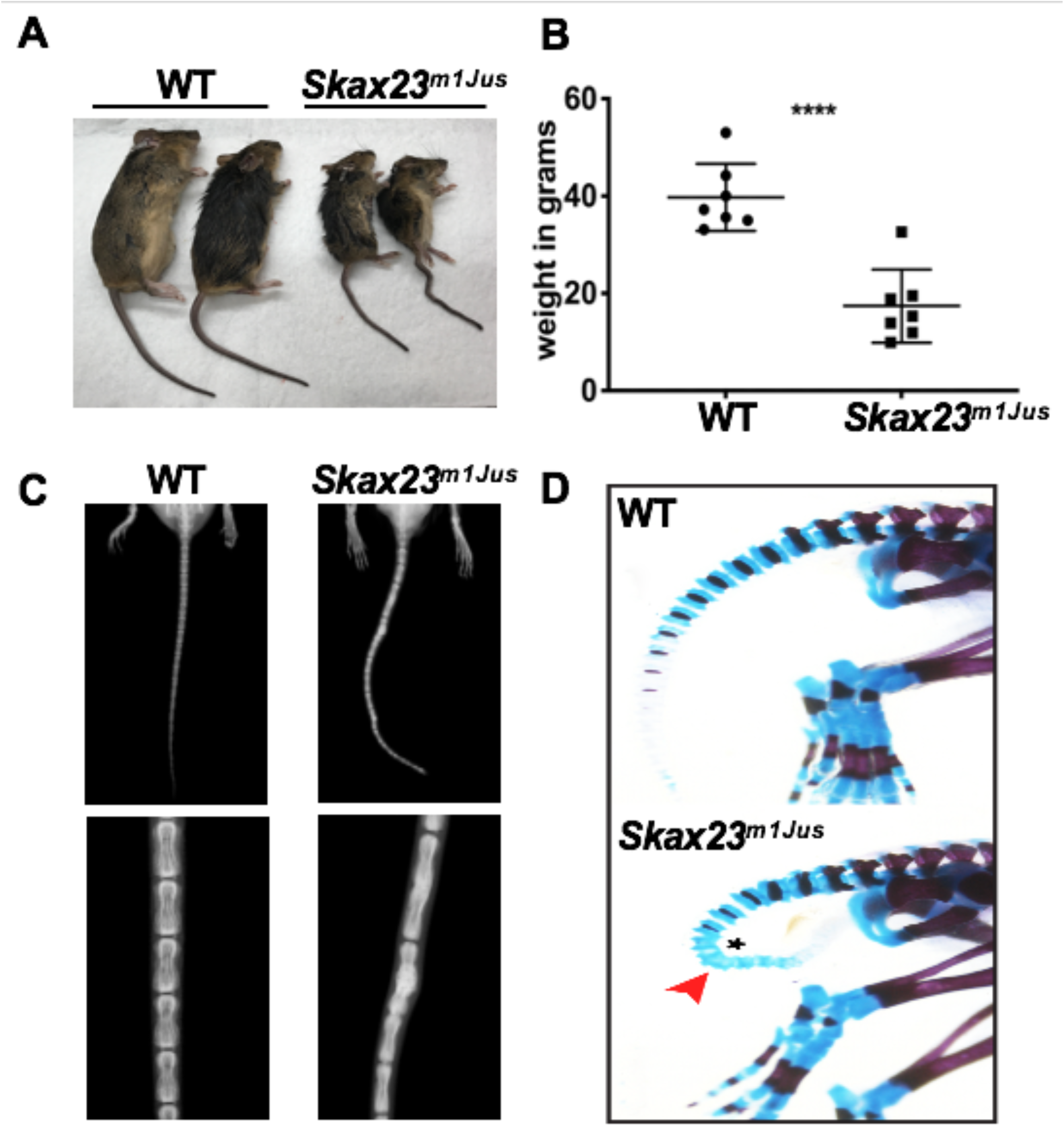
*Skax23*^*m1Jus*^ mice grow poorly and exhibit a kinky tail defect due to delayed endochondral ossification. (A) The *Skax23*^*m1Jus*^ mouse mutant was generated and identified as part of the chromosome 11 balancer mutagenesis screens at the Mouse Mutagenesis and Phenotyping Center for Developmental Defects at Baylor College of Medicine in Houston (http://www.mouse-genome.bcm.tmc.edu). *Skax23*^*m1Jus*^ was identified as a dominant mutant with a kinky tail phenotype and poor growth. (B) Mutant adult mice (n=7) were significantly smaller than wildtype (n=7). (C) X-ray analysis of adult mice show abnormal fusion and hypertrophy of tail bone joints in *Skax23*^*m1Jus*^ mice when compared with wild type littermates (low power view 1.5-2x, high power view 3x). (D) Whole-mount staining of newborn mice using Alcian blue (for cartilage) or Alizarin red (for bone) was performed. *Skax23*^*m1Jus*^ newborn mice had kinky tails (red arrowhead) with delayed endochondral ossification (asterisk).

### Genetic mapping

For genetic mapping studies, a total of 21 mice with a kinked tail phenotype from 4 backcrosses (4 N1F1, 5 N2F1, 4 N3F1 and 8 N4F1) were genotyped with 180 SNPs distributed randomly across the genome. These markers were informative for both C57BL/6J and 129S6/SvEv strains and the average intermarker distance was 15 Mb. Of these, 26 markers failed to genotype and 5 were not informative in both strains resulting in 149 markers for genotyping data analyses. Candidate regions were prioritized by the percentage of C57BL/6J heterozygosity in affected mice. Fine mapping of the candidate region on chromosome 5 was done by genotyping 20 affected mice included in the original whole genome scan (4 N1F1, 5 N2F1, 3 N3F1 and 8 N4F1) and 10 affected mice obtained from a 5^th^ backcross to 129S6/SvEv animals (10 N5F1) with 14 SNPs selected from within this region.

### Whole exome sequencing

Whole exome sequencing (WES) was conducted in 2 affected mice (one N1F1 and one N1F4) and 1 unaffected 129S6/SvEv mouse. WES capture was done using the Sureselect Mouse All Exon 50 Mb (Agilent technologies). Paired-end sequencing was performed on an AB Hiseq 2000. Raw data were aligned to the C57BL/6 NCBI37/mm9 genome reference build using the Burrows-Wheeler Aligner. Duplicate reads were removed, remaining reads were realigned locally, and variants were called using GATK 2.6.4. Variant annotation was done using Annovar (http://www.openbioinformatics.org/annovar/). This resulted in variant coverage of 15X average. Mutations mapping to the candidate region on chromosome 5 that were heterozygous in both affected mice and absent from the C57BL/6 reference genome and from the 129S6/SvEv mice were prioritized for validation by Sanger sequencing. Genotyping protocols and PCR primers are described in detail in supplemental methods.

### Cell culture

Spleens were isolated, crushed to generate a single cell suspension and strained, followed by red cell lysis performed using ACK Lysing Buffer (Lonza). Total spleen cells were washed, resuspended in complete media (RPMI, 10% FBS, 2 mM L-glutamine, 1% Pen/Strep, 50 μM 2-mercaptoethanol, 10 mM HEPES) at a concentration of 2×10^6^ cells/ml. LPS was added to culture media at 10 μg/ml for 24-72 hours. Cell number and viability were assessed using an ADAM MC Auto Cell Counter.

### Western blot

Total spleen cells were harvested at 48 hours of culture and protein extracts were prepared for western blots as previously described, using antibodies listed in Supplemental Materials, Table S1^37^.

### Polysome profiling

Total spleen cells grown in complete media supplemented with LPS (10 μg/ml) for 36 hours were processed for polysome profiling as previously described^31^.

### Statistics

The Student’s t-test in GraphPad Prism^®^ 9 was used to determine statistical significance. (* p<0.05; ** p<0.01, *** p<0.001, **** p<0.0001 for all figures). Error bars were determined using the standard error of the mean. Statistics for Mendelian ratios were calculated using Chi squared (χ^2^) analysis.

The primary data are available on request (S.A.S.).

## Results

### Description of the *Skax23*^*m1Jus*^ kinky tail dominant mutant

*Skax23*^*m1Jus*^ mutant mice can be easily identified at birth due to their small size and kinked tail (Fig. 1A, sFig1). Mice that survive the neonatal period continue to grow poorly and surviving adult mutant mice are about half the weight of wildtype mice (Fig. 1B). Examination of the skeleton via x-ray of adult mice showed abnormal hypertrophy and fusion of tail bone joints (Fig. 1C). Additional skeletal analysis of newborn *Skax23*^*m1Jus*^ mice demonstrated reduced skulls and bone lengths although only femur lengths were significantly decreased (sFig. 1B–C). However, all newborn *Skax23*^*m1Jus*^ mice exhibited kinky tails with no evidence of endochondral ossification in the lower caudal vertebrae (Fig. 1D).

### Identification of *Rpl5 (uL18)* as the gene defective in *Skax23*^*m1Jus*^

To map the genetic defect in *Skax23*^*m1Jus*^/+ mice, mice with a kinky tail phenotype were subjected to 5 consecutive backcrosses to 129S6/SvEvTac animals. A whole genome scan using 149 SNPs informative for both C57BL/6J and 129S6/SvEv was conducted on 21 N1F1-N4F1 mice with a kinky tail phenotype. Since the ENU-induced mutation was generated on the C57BL/6J background, regions with the highest percentage of heterozygosity for C57BL/6J were prioritized. This analysis identified a ~ 41 Mb region on chromosome 5 that had the highest % heterozygosity in C57BL/6J animals (sFig. 2A). Analysis of 13 SNPs from the whole genome scan that covered this region identified a haplotype consisting of 3 SNPs (rs13478451, rs32103915 and rs29534493) that was heterozygous in the C57BL/6J genome in all 21 mice analyzed (sFig. 2B). Fine mapping within this candidate region was done by genotyping 20 affected mice included in the original whole genome scan and 10 additional affected mice obtained from a 5^th^ backcross with 14 SNPs selected from this region. This fine mapping has reduced this region to 11 Mb from 107112543bp (rs13478444) to 118405516bp (rs13478483) on the NCBI37/mm9 assembly (sFig. 2C).

**Figure 2:**
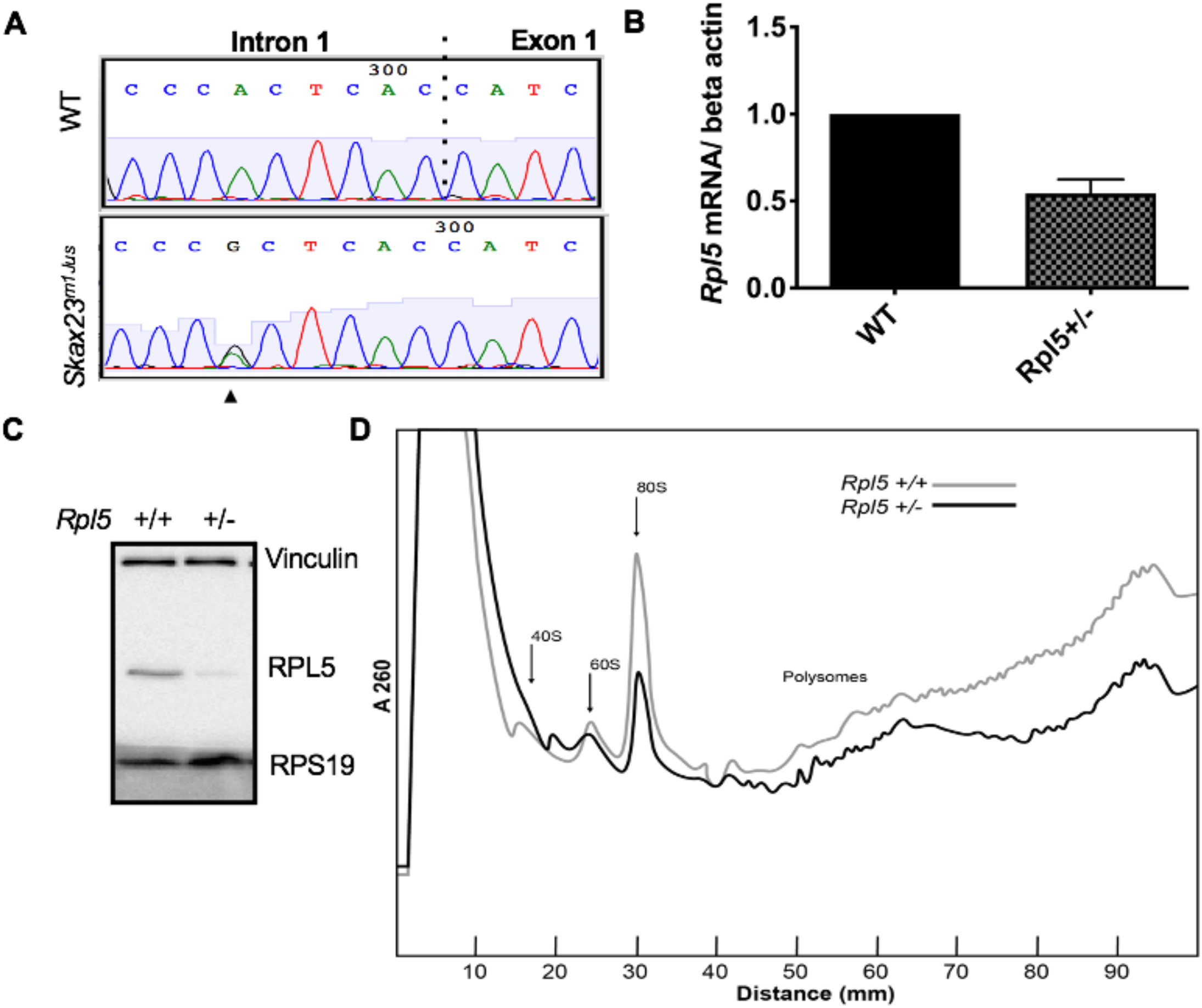
*Rpl5* intronic mutation identified in *Skax23*^*m1Jus*^ mice leads to decreased mRNA, protein levels and polysome defects. A. Chromatogram of reverse strand partial sequences of exon 1 and intron 1 of *Rpl5* showing the heterozygous mutation (indicated by triangle) at the 6^th^ nucleotide in intron 1. The exon-intron junction is indicated by a dotted line. B. RT-qPCR showed an approximately 50% reduction in *Rpl5* mRNA in the *Rpl5+/−* mice when compared to wild type. C. Western blot using vinculin and RPS19 as loading controls showed reduced levels of RPL5 protein in *Rpl5+/−* mice when compared to WT. D. Results from 3 separate experiments indicate *Rpl5*+/− mice have reduced 60S subunits, 80S ribosomes and polysomes, consistent with *Rpl5* haploinsufficiency.

To identify the gene defective in *Skax23*^*m1Jus*^ mice, we conducted whole exome sequencing (WES) on 2 affected and 1 unaffected 129S6/SvEv mouse. Mutations present within the candidate region on chromosome 5 in the 2 affected mice and absent from both parental strains and the reference genome were prioritized for study. A total of 4 mutations obeying these criteria were identified (data not shown). Following validation by Sanger sequencing, only one intronic mutation in *Rpl5* (aka *uL18*), c.3+6C>T, was confirmed (Fig. 2A)^20^. The *RPL5* gene is mutated in some forms of Diamond-Blackfan anemia (DBA) and a different ribosomal protein mutant (*Rps7* or *eS7*) found in DBA patients produces a kinked tail and a smaller size in another mouse model, making this mutation an excellent candidate^38^. For genotype-phenotype studies, genotyping was done on a total of 228 mice with a straight tail and 95 mice with a kinky tail. The c.3+6C>T heterozygous mutation was absent from all mice with a straight tail and segregated with the kinky tail phenotype in all 95 mice analyzed, indicating complete penetrance. No homozygous mice bearing this variant were identified, suggesting embryonic lethality in the homozygous state. This variant was also absent in 28 other inbred strains (data not shown), confirming that it is not a rare variant and that it is specific to the phenotype.

### The c.3+6C>T intronic mutation results in *Rpl5* haploinsufficiency

RT-qPCR analysis demonstrated an approximately 50% reduction in *Rpl5* mRNA in *Skax23*^*m1Jus*^ +/−mutants compared with WT mice (Fig. 2B). Western blot analysis (in triplicate) showed reduced RPL5 protein in mutant mice (Fig. 2C). Analysis of 3 separate polysome experiments demonstrated reduced 60S subunit, 80S ribosomes and polysomes, consistent with *Rpl5* haploinsufficiency (Fig. 2D). We will refer to the *Rpl5*^*Skax23-Jus*^ mutation as *Rpl5*+/− throughout the manuscript.

### Neonatal *Rpl5*+/− mice have a macrocytic anemia, cardiac defect and increased mortality

Newborn mice from WT x *Rpl5+/−* matings were genotyped (14 litters, n=58). The percentage of *Rpl5+/−* offspring was significantly lower than expected by Mendelian predictions (Table 1A). Three litters were followed over time to determine the percentage of newborn pups that died after birth and prior to weaning, (WT n=8, *Rpl5+/−* n=5). Three mutant pups (60%) died postnatally, 2 on day 1 and 1 on day 21 (Table 1B). The pup that died 21 days after birth was found to have severe pancytopenia (WBC 1 × 10^9^/L, hemoglobin (Hgb) 3.5g/dL, platelets 11,000 × 10^9^/L). Two pups from additional litters were runted and appeared lethargic and therefore had to be euthanized 14-15 days postnatally. Complete blood count (CBC) analyses were performed at the time of euthanasia and demonstrated Hgbs 6.6 and 7.3g/dL, which were significantly lower than the Hgb of their WT littermate control (Hgb 14.7g/dL). One of the 2 latter pups euthanized was also thrombocytopenic with a platelet count of 14,000, while the other had a normal platelet count.

**Table 1:** Mendelian ratio of pups at birth. (A) WT x *Rpl5+/−* matings were performed and 14 litters were genotyped (n=58). The percentage of *Rpl5+/−* offspring was significantly lower than the Mendelian prediction. χ^2^ analysis, p=0.0039. (B) 3 litters were followed to determine the % of newborn pups that died after birth and prior to weaning, WT n=8, *Rpl5+/−* n=5. 3/5 mutant pups (60%) died (2 pups on day 1 and 1 pup at 21 days) during the stated period.

|  | *Rpl5*<sup>+/+</sup> | *Rpl5*<sup>+/-</sup> | Total | % *Rpl5*<sup>+/-</sup> |
| --- | --- | --- | --- | --- |
| Observed | 40 | 18 | 58 | 31 |
| Predicted | 29 | 29 | 58 | 50 |

**Table 1**
|  | *Rpl5*<sup>+/+</sup> | *Rpl5*<sup>+/-</sup> | % *Rpl5*<sup>+/-</sup><br>*death* |
| --- | --- | --- | --- |
| Died before wean | 0/8 | 3/5 | 60% |

We next analyzed CBCs of newborn mice at days 1-3 postnatally (WT n=10 and *Rpl5+/−* n=7) and found that *Rpl5+/−* mice had macrocytic anemia (Figs. 3A–B) but normal WBC and platelet counts (Figs. 3C–D). Newborn *Rpl5+/−* livers had normal cellularity (Fig 3E) but cell suspensions from these livers appeared paler compared with those of WT livers. Additionally, the Ter119 mean fluorescent intensity, as determined by flow cytometry, was reduced in *Rpl5+/−* compared to WT livers (Figs. 3F–G). Further flow analysis of erythroid differentiation showed no significant difference in the proerythroblast (CD44^hi^ Ter119^low^) population between mutant and WT livers (WT n=3, *Rpl5+/−* n= 3) (Figs. 3H–I)^39^.

**Figure 3:**
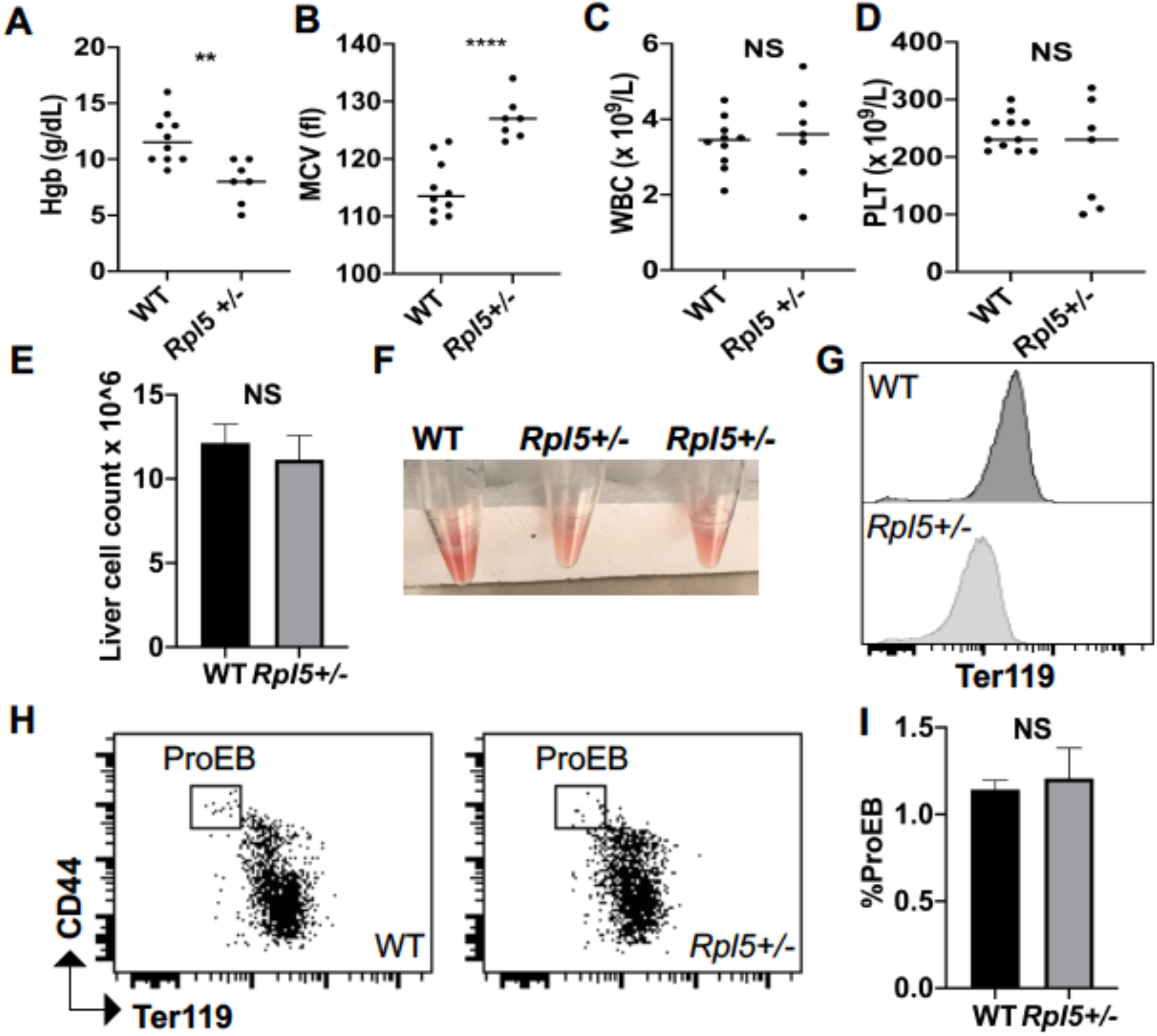
*Rpl5*+/− newborn mice have macrocytic anemia. Newborn mice (days 1-3) were euthanized and blood counts analyzed (WT n=10, *Rpl5+/−* n= 7). *Rpl5+/−* mice exhibited evidence of anemia (A) with significantly lower hemoglobin (Hgb) values and (B) macrocytosis as demonstrated by an elevated MCV. There was no significant difference in white blood cells (WBC) (C) or platelet (PLT) counts (D) between *Rpl5+/−* and WT mice. The line represents the median value for each group. Newborn livers (day 1-2) were isolated and flow cytometry was performed to analyze erythroid differentiation. (E) There was no difference in liver cellularity but all *Rpl5*+/− mice had paler livers (F) with decreased Ter119 mean fluorescent intensity (G). (H, I) There was no difference in proerythroblast numbers (WT n=3, *Rpl5+/−* n= 3).

Newborn mice were also analyzed for the presence of craniofacial abnormalities (common in DBA patients with heterozygous *RPL5* mutations) and cardiac defects. Histologically, out of 6 *Rpl5+/−* mice evaluated, 5 mice showed evidence of ventricular septal defect (VSD) and 1 mouse had double outlet right ventricle in addition to VSD (sFig. 3A, DORV not shown). There was no evidence of VSD by histology in surviving adult *Rpl5+/−* mice (WT n=2 and mutant n=7). These results suggest that presence of a VSD was associated with increased risk of death of *Rpl5+/−* newborn animals. There was no evidence of a craniofacial defect by micro-CT or histology in *Rpl5*+/− mice (sFigs. 3B–C)^40^.

### *Rpl5*+/− mice exhibit a severe erythroid maturation defect at E12.5

In order to determine whether the onset of the anemia in *Rpl5*+/− mice was pre-or postnatal, we analyzed E12.5 and E14.5 embryos generated from timed matings between WT females x *Rpl5+/−* males. Genotyping of 5 litters at E12.5 (n=38) revealed 28 live embryos with no significant difference between observed and expected genotypes (Table 2A). However, there were 10 dead embryos observed at different stages of reabsorption, all of which had the *Rpl5*+/− genotype, indicating that death in some mutants occurred prior to E12.5 (Table 2B). E12.5 *Rpl5+/−* embryos were significantly growth retarded and pale, with 2 mutant embryos showing signs of intracranial bleeding (Fig. 4A; sFig. 4, embryos #20-5, #20-6). Dead embryos were found at both E12.5 (sFig. 4B, #3-5) and E14.5 (sFig. 5, #19-3). At E12.5, compared to WT fetal livers (FL), *Rpl5+/−* FL had a significantly lower number of total live cells and notably, a smaller proportion of Ter119+ cells. (Fig. 4B–C). Mutant embryos at E14.5 had less prominent growth retardation and pallor compared to E12.5 *Rpl5+/−* embryos (Fig. 4D, sFig. 5). At E14.5, the FL cellularity remained significantly diminished in *Rpl5+/−* embryos; however the percentage of Ter119 positive cells had clearly increased (compared to E12.5), albeit it remained lower than normal (Figs. 4E–F).

**Table 2:**
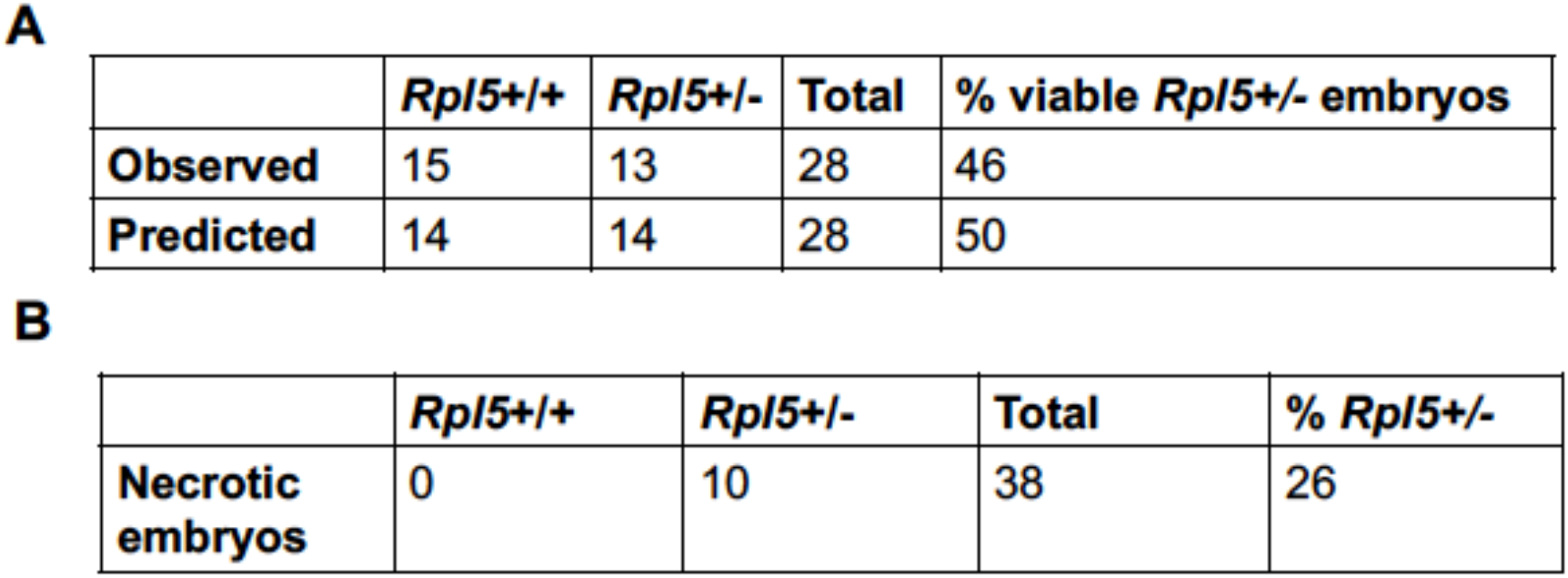
Genotyping of embryos at E12.5. Timed matings were set up with WT females x *Rpl5+/−* male animals. Pregnant females were euthanized at E12.5 and genotyping was performed on all embryos. (A) Genotyping performed on 5 litters (n=38) revealed 28 live embryos with no significant difference in observed to expected Mendelian ratio (χ^2^ analysis, p=0.7). (B) An additional 10 *Rpl5+/−* embryos (26%) were found to be dead or necrotic (dead embryos not shown in A).

|  | *Rpl5*<sup>+/+</sup> | *Rpl5*<sup>+/-</sup> | Total | % viable *Rpl5*<sup>+/-</sup> embryos |
| --- | --- | --- | --- | --- |
| Observed | 15 | 13 | 28 | 46 |
| Predicted | 14 | 14 | 28 | 50 |

**Table 2**
|  | *Rpl5*<sup>+/+</sup> | *Rpl5*<sup>+/-</sup> | Total | % *Rpl5*<sup>+/-</sup> |
| --- | --- | --- | --- | --- |
| Necrotic embryos | 0 | 10 | 38 | 26 |

**Figure 4:**
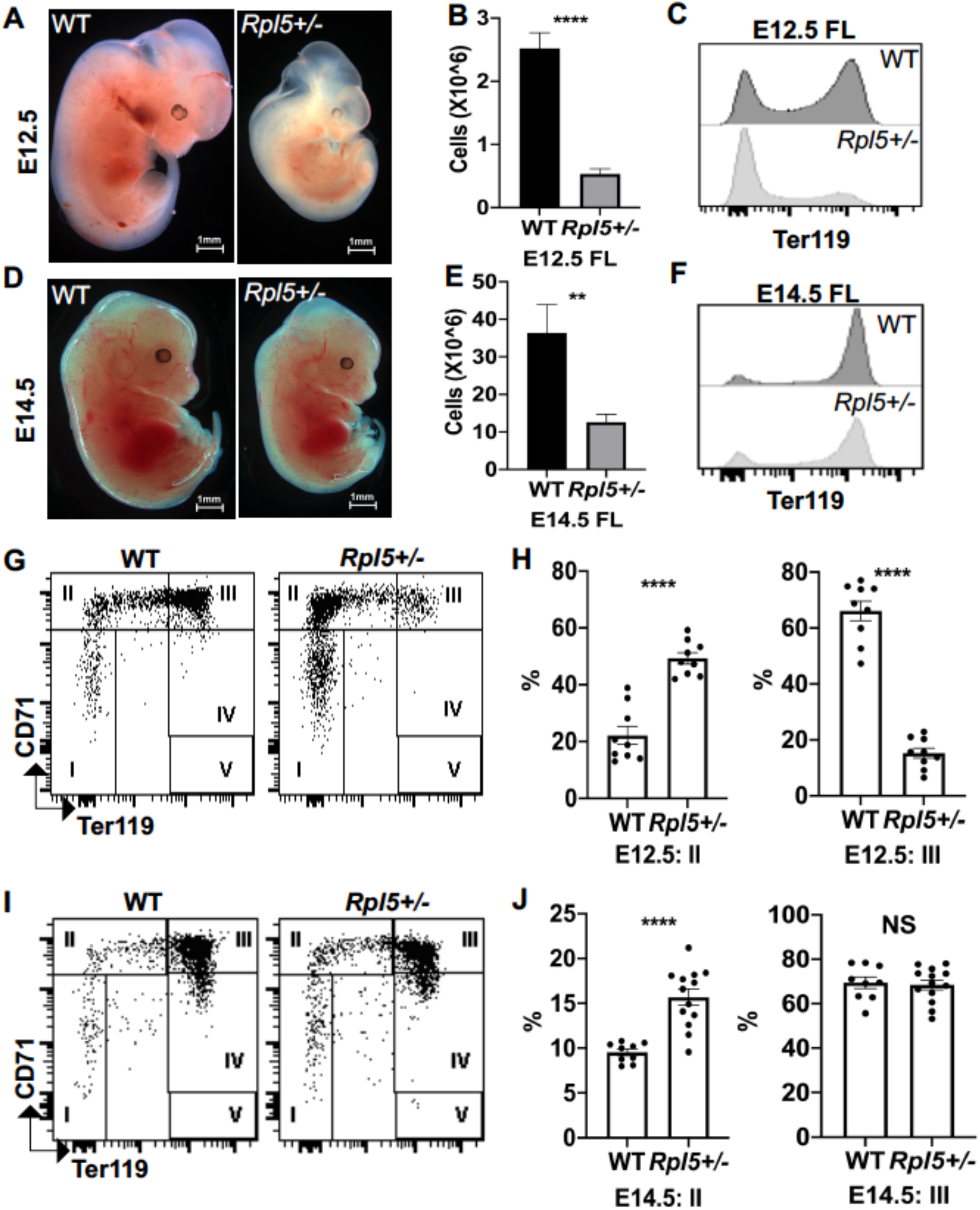
*Rpl5+/−* mice have a severe block in erythroid differentiation at E12.5 that improves by E14.5. (A) Timed WT x *Rpl5+/−* matings were set up to collect E12.5 and E14.5 embryos. E12.5 *Rpl5+/−* embryos were generally growth retarded and pale. (B) Mutant E12.5 embryos displayed a marked reduction in fetal liver (FL) cellularity with fewer FL Ter119+ cells (C). (D) E14.5 mutant embryos had improved growth, pallor, FL cellularity (E) and increased Ter119+ cells (F). (G) Analysis of red cell differentiation by CD71 and Ter119 showed a significant delay in the transition from population II to III in the FL of *Rpl5+/−* mice at E12.5 (quantified in H). (I) A similar analysis conducted on E14.5 FL cells showed an improvement in the E12.5 differentiation block with no difference in population III at this stage (J). E12.5 WT n=9, *Rpl5+/−* n=9 (3 separate litters); E14.5 WT n=9, *Rpl5+/−* n=12 (4 separate litters).

Further analysis of FL erythropoiesis was performed at E12.5 and E14.5 by examining CD71 and Ter119 expression^41,42^. At E12.5, erythroid cells at stage II of differentiation (CD71+/Ter119-/low) were increased in *Rpl5+/−* FL while erythroid cells at stage III of differentiation (CD71+/Ter119+) were reduced, suggesting a delay in erythroid differentiation resulting from *Rpl5* haploinsufficiency (Figs. 4G–H). Additional analysis using CD44 and Ter119 showed a significant (3-fold) increase in the proerythroblast (proEB) population in mutant animals, again indicative of an erythroid differentiation block (sFigs. 6A-B; E12.5 WT n=9, *Rpl5+/−* n=9 from 3 litters). A similar analysis of E14.5 FL cells showed an improvement in the erythroid differentiation block, with no difference in the percentage of erythroid cells at stage III of differentiation between *Rpl5+/−* and WT FL (Figs. 4I–J). Additional analysis demonstrated that the proportion of proEB in the E14.5 livers remained elevated (albeit was improved) at 1.5-fold compared to WT livers (sFigs. 6C-D; E14.5 WT n=9, *Rpl5+/−* n=12; 4 litters).

E12.5 FL cells were next analyzed to assess whether there was evidence for increased apoptosis of *Rpl5+/−* erythroid cells^43^. Populations II and III were analyzed for apoptosis by staining Annexin V and DAPI (sFig. 7). There was no significant difference in the percentage of population II or III erythroid cells undergoing early or late apoptosis between *Rpl5+/−* and WT embryos (WT n=6, *Rpl5+/−* n=6; 2 litters).

### Adult *Rpl5*+/− mice exhibit no anemia or hematopoietic defects

Analysis of *Rpl5*+/− mice at 7 weeks of age showed no evidence of anemia or blood count abnormalities, when compared with WT controls (sFigs. 8A-F; WT n=10, *Rpl5+/−* n=10). A similar analysis of aged mice also did not demonstrate any blood count abnormality (sFigs. 8G-L; WT n= 11, *Rpl5+/−* n=7). Bone marrows (BM) of 5-month-old mice were analyzed by flow cytometry for red cell markers (Ter119, CD44) after exclusion of nonerythroid cells (CD45+, Gr-1, CD11b, CD3) and dead cells. No significant difference in Ter119 mean fluorescence intensity was found between the BM of *Rpl5+/−* and WT animals (sFig. 9A). There was also no significant difference in percentage of the proerythroblast population between *Rpl5+/−* and WT animals (sFigs. 9B-C; WT n=3, *Rpl5+/−* n= 3). Further flow analysis of the bone marrows of adult mice did not reveal significant differences in BM cellularity and the percentage of early stem/progenitor cells, mature myeloid cells, or mature lymphocyte subpopulations (sFig. 10)^44,45^.

## Discussion

In this study, we identified and characterized a novel intronic ENU-induced mutation (*Skax23*^*m1Jus*^) in the *Rpl5* gene, affecting the 6^th^ nucleotide in intron 1 (c.3+6C>T mutation). This mutation maps to the candidate region on chromosome 5 and segregates with the kinky tail phenotype with 100% penetrance. It was absent from the parental strains and 28 other inbred strains suggesting it was the cause of the phenotype. Our best hypothesis at present is that the c.3+6C>T mutation affects splicing, possibly causing intron 1 retention and leading to nonsense-mediated mRNA decay. Alternatively, it might affect intronic transcription regulatory elements of an unknown nature. Additional *in vitro* studies are needed to elucidate the mechanism by which this mutation affects *Rpl5* mRNA and protein expression levels. This mutation leads to a stable decrease in *Rpl5* mRNA and protein levels and to polysome defects making it distinct from the other published *Rpl5* zebrafish and mouse models. The morphological defects (small size and kinky tail) in this model mouse are similar to the previously described *Rps7* mutant mouse and correspond to the increased incidence of congenital abnormalities in patients with *RPL5* variants^7,28,29,38^.

Our initial characterization of adult *Rpl5+/−* mice demonstrated a surprising lack of anemia, without detectable defects in stem and progenitor cells. However, the observation of increased mortality of newborn mutant pups led to the discovery of a macrocytic anemia at birth that worsened in some pups, possibly explaining their demise. However, the anemia observed in *Rpl5+/−* mice at birth had completely resolved in adult animals that survived to 7 weeks of age. Examination of FL cells revealed a more severe erythroid differentiation defect at E12.5 that was slowly resolving by E14.5 in surviving animals. These variable observations seen at different developmental time points in *Rpl5+/−* mice were reminiscent of the variable anemia and spontaneous unexplained remissions that can occur in some human DBA patients. The mechanism underlying this phenomenon is unknown but environmental and/or cell-extrinsic factors such as elevated reactive oxygen species have been proposed to play a role^9^. As the inbred mice examined in this study have been backcrossed >7 times with no change in phenotype (now >98% 129/SvJ congenic by Jax genome scan), genetic polymorphisms are unlikely to influence this variably penetrant anemia. The mice are also housed in a specific pathogen free environment and monitored frequently for invasive pathogens so it is equally unlikely that viral or other infections have contributed to the anemia. Furthermore, as the variation in presentation occurs within the same litter in the same mouse cage (i.e. some mutants succumb prior to weaning while other mutant littermates survive to adulthood); the variability in phenotype is unlikely to be due to major environmental changes, but this certainly requires further study. In our current work, we have not examined if somatic reversion of *Rpl5* occurs and leads to selection of more fit clones that can rescue the hematopoietic defect, which has been described previously in both DBA and Fanconi anemia^46–50^.

A question raised by this DBA model is whether the variable anemia phenotype is strain-dependent. Our mice are currently maintained on a 129/SvJ background but we have not yet studied this erythroid phenotype in other common inbred backgrounds such as C57BL/6. Also, it is unclear whether the variable anemia observed in our model is also present in other DBA animal models resulting from mutations in other DBA genes or if this phenotype is specific to *Rpl5* or to this particular *Rpl5* intronic mutation. Of note, variants in noncoding and splice site regions of genes encoding RPs have previously been reported in DBA patients^7,51,52^. As the variation in phenotype is present in DBA patients of all genotypes, it is unlikely that the findings reported here are specific for *Rpl5*, but this requires further study.

The non-erythroid (skeletal or cardiac) manifestations of DBA have not been as extensively studied as the erythroid phenotype. Here, we show that the kinky tail in *Rpl5+/−* mutant mice is due to delayed endochondral ossification. Why RP haploinsufficiency only affects certain tissues and not others is an area that requires further study but is possibly due to selective decreased translation of vulnerable mRNAs. In a previous study describing a *Rpl38* haploinsufficient mouse, skeletal defects were concluded to be due to decreased translation of specific *Hox* mRNAs, while global translation appeared unaffected^53^. As the *RPL5* genotype is strongly associated with craniofacial malformations, we initially extensively analyzed mice for such defects that could result in poor feeding and growth retardation. However, we were unable to find conclusive evidence for craniofacial malformations by histology or by micro-CT. The presence of the cardiac defects (including VSD) was not surprising, as a subset of DBA patients also have such defects. In addition, neural crest cells have been shown to play a major role both in both craniofacial and cardiac morphogenesis and neural crest defects can lead to septal defects like VSD^54–56^. Moreover, several genes involved in ribosome biogenesis are required for neural crest cell proliferation and survival, and defects in these genes result in severe craniofacial defects^57^.

The observation reported here that most newborn *Rpl5+/−* mice have VSDs, while surviving mice do not bears some consideration. It is possible that the cardiac progenitor cells are subject to the same process that creates a range of severity in the erythroid lineage. Therefore, the presence of a VSD may be a way to track those mice that are on a trajectory for total erythroid failure. The finding of variable thrombocytopenia is consistent with the phenotype in DBA patients and also warrants further study^58,59^. We therefore propose that this *Rpl5*^*ska23-Jus*^ mutant mouse model can be used to study the underlying mechanisms leading to variable penetrance and remission of anemia in DBA.

Here we described a murine model of DBA that appears to recapitulate several features of the human disease that were not found in other previously described mouse models, including the variable anemia with spontaneous remission. To date, to our knowledge, no other animal models have captured this dynamic erythroid presentation, which is a key feature of human DBA. We also propose using VSD as a possible way to track those mice whose trajectory might lead to a more severe erythroid defect so that the underlying mechanism(s) might be more fully explored. This report suggests that the variation in penetrance in DBA may be stochastic without the influence of genetic modifiers or significant environmental influences. Future studies aimed at improving our understanding of the mechanism of this variable penetrance may reveal novel biological pathways relevant to RP deficiency states. This novel *Rpl5*^*ska23-Jus*^ mutant strain presents a unique opportunity to study fundamental mechanisms involved in normal and disordered erythropoiesis and should be useful in testing new therapeutic approaches.

## Acknowledgments

The authors thank Michael Clemente in the core flow cytometry facility at Western Michigan University School of Medicine, the members of the vivarium at Western Michigan University School of Medicine and the University of Michigan, the Physiology Phenotyping Core at University of Michigan, Evan Keller for the use of x-ray equipment, Jinlu Dai for assistance with imaging, Kim-Chew Lim, Sowmya Balasubramanian, Mark Chiang, Jordan Shavit, Emily Philipp-Petrick, Selest Nashef, Stephan Owens and the members of the Rothstein laboratories for their support and assistance.

This work was supported by M-cubed, a research seed funding program for faculty at the University of Michigan (S.A.S., V.K., R.F., R.K.), Canadian Institutes of Health (Z.K.), National Institute of Health Grant R01 HL148333 (R.K.), U01 HD39372 (M.J.J.), and RO1 CA115503 (M.J.J.), The University of Michigan Rogel Cancer Center P30CA046592 grant (R.K.), T32HL007622 (M.J.).

## Authorship

Contribution: F.M., G.S., C.P., E.S. performed the experiments, interpreted the data and edited the manuscript; L.Y., P.L., A.L., M.G., M.J., M.J.J. designed and performed the experiments, interpreted the data and edited the manuscript; Y.W., G.M., R.K., Q.L, R.F., T.L.R., J.D.E., V.K. designed the experiments, interpreted the data, and edited the manuscript; Z.K. and S.A.S. designed and performed the experiments, interpreted the data, and wrote and edited the manuscript.

