## Supplemental Figures for "A new murine *Rpl5* (*uL18*) mutation provides a unique model of variably penetrant Diamond Blackfan Anemia"

**Table S1:** Antibodies/reagents used for western blot/erythroid studies

| **Antibody** | **Source** | **Product number** | **Experiment** |
|---|---|---|---|
| mouse monoclonal anti-vinculin, clone hVIN-1 | Sigma | V9131 | Western blot 1:10,000 |
| rabbit monoclonal anti-RPS19 | Abcam | ab181365 | Western blot 1:10,000 |
| rabbit polyclonal anti-RPL5 | Cell Signaling | 14568 | Western blot 1:1000 |
| APC anti-mouse Ter119 | Biolegend | 116212 | Flow cytometry E12.5, E14.5 |
| FITC anti-mouse CD71 | Biolegend | 113806 | Flow cytometry E12.5, E14.5 |
| APC/Cy7 anti-mouse CD44 | Biolegend | 103028 | Flow cytometry E12.5, E14.5 |
| PE Annexin V | BD Biosciences | 556421 | Apoptosis flow cytometry E12.5 |
| DAPI | Biolegend | 422801 | Flow cytometry E12.5, E14.5 |
| LIVE/DEAD™ Fixable Aqua stain | ThermoFisher Scientific | L34957 | Newborn flow cytometry |
| PE CD45 | BD Biosciences | 553081 | Newborn flow cytometry |
| PE Gr-1 | BD Biosciences | 553128 | Newborn flow cytometry |
| PE CD11b | BD Biosciences | 557397 | Newborn flow cytometry |
| PE CD3 | BD Biosciences | 555275 | Newborn flow cytometry |
| FITC Ter119 | BD Biosciences | 561032 | Newborn flow cytometry |
| APC CD44 | BD Biosciences | 559250 | Newborn flow cytometry |

**Table S2:** Antibodies used for flow cytometry studies of stem/progenitor, mature myeloid/lymphoid cells

| Target | Fluorochromes | Clone | Source |
|---|---|---|---|
| CD150 | PE-Cy7 | TC15-12F12.2 | Biolegend |
| CD48 | AlexaFluor700 | HM48-1 | Biolegend |
| CD16/32 | PE-Cy7 | 93 | Biolegend |
| CD34 | FITC | RAM34 | eBiosciences |
| Sca-1 | Percp-Cy5.5 | D7 | Biolegend |
| cKit | APC | 2B8 | Biolegend |
| Ter119 | PE,FITC | TER-119 | Biolegend |
| B220 | PE,FITC | RA3-6B2 | Biolegend |
| CD2 | PE,FITC | RM2-5 | Biolegend |
| CD3 | PE,FITC | 17A2 | Biolegend |
| CD5 | PE,FITC | 53-7.3 | Biolegend |
| CD8 | PE,FITC | 53-6.7 | Biolegend |
| CD11b | APC | M1/70 | Biolegend |
| Gr1 | PE-Cy7, PE, FITC | RB6-8C5 | Biolegend |

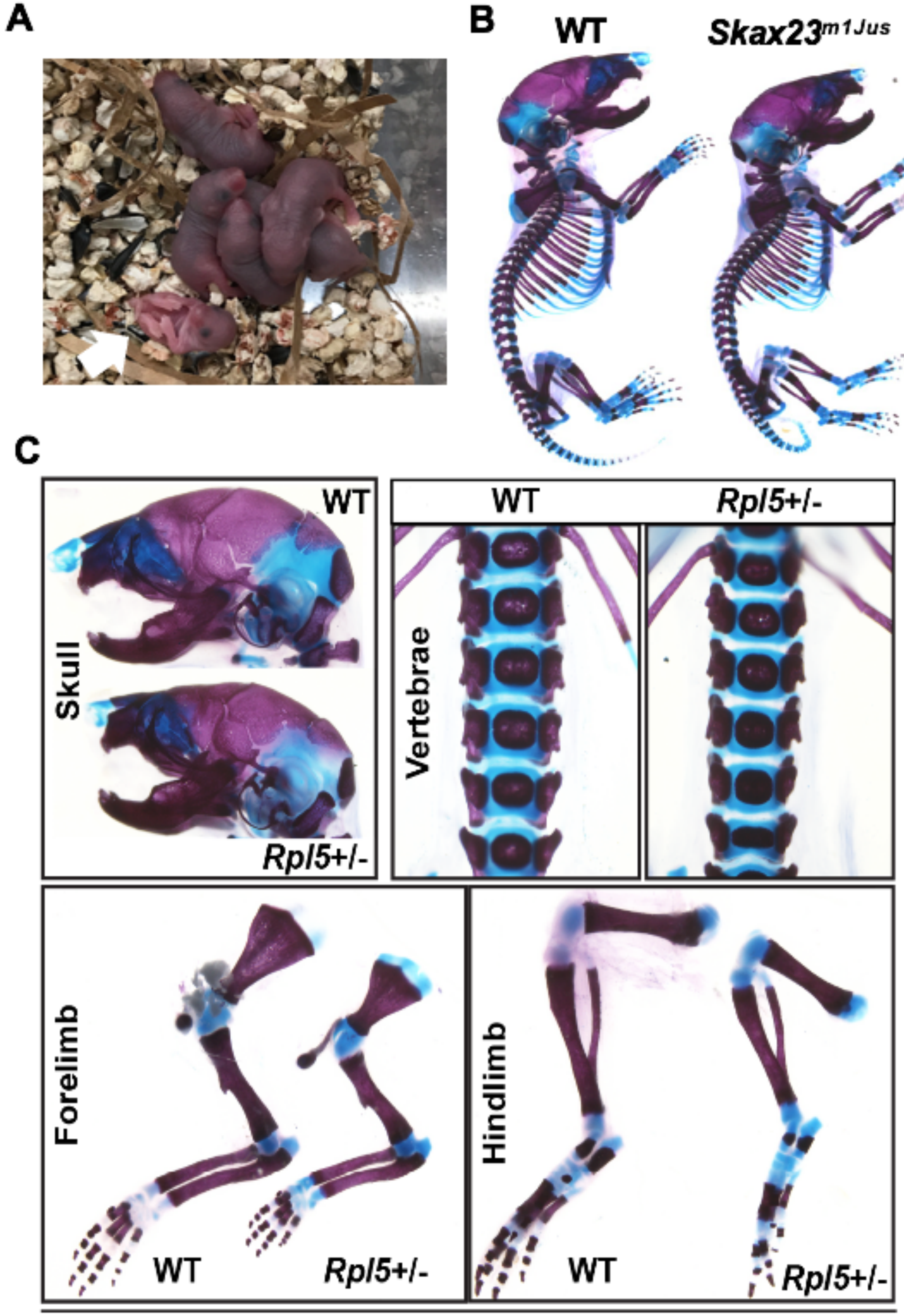


**Supplemental figure 1**

A

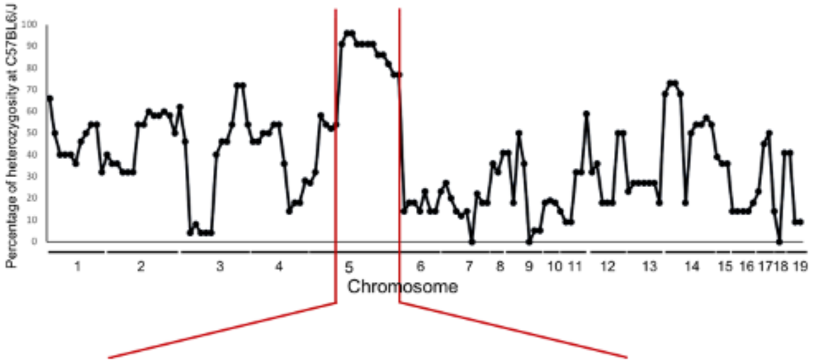


B

| SNP | Position | 129S6/SvEv | C57BL/6 | Mice with kinky tail phenotype | | | | | |
|---|---|---|---|---|---|---|---|---|---|
| | | | | N=11 | N=6 | N=1 | N=1 | N=1 | N=1 |
| rs13478433 | 104357881 | TT | CC | CT | TT | TT | CT | TT | TT |
| rs13478451 | 109083680 | CC | TT | CT | CT | CT | CT | CT | CT |
| rs32103915 | 112501418 | AA | TT | AT | AT | AT | AT | AT | AT |
| rs29534493 | 115004548 | TT | CC | CT | CT | CT | CT | CT | CT |
| rs13478483 | 118405516 | TT | CC | CT | CT | CT | CT | CT | TT |
| rs8239888 | 122017784 | CC | TT | CT | CT | CT | CT | CT | CC |
| rs3671202 | 125290118 | CC | TT | CT | CT | CT | CT | CT | CC |
| rs13478518 | 128264874 | CC | TT | CT | CT | CT | CT | CT | CC |
| rs13478539 | 133538654 | TT | CC | CT | CT | CT | CT | TT | TT |
| rs13478542 | 135358216 | CC | TT | CT | CT | CT | CT | CC | CC |
| rs29779884 | 141325735 | TT | CC | CT | CT | CT | TT | TT | TT |
| rs13478574 | 144868252 | CC | TT | CT | CT | CC | CC | CC | CC |
| rs3718776 | 150393126 | TT | CC | CT | CT | TT | TT | TT | TT |

C

| SNP | Position in MB | 129S6/SvEv | C57BL/6 | Mice with kinky tail phenotype | | | |
|---|---|---|---|---|---|---|---|
| | | | | N= 12 | N=1 | N=10 | N=7 |
| rs13478433 | 104.357881 | TT | CC | CT | TT | TT | TT |
| rs29559613 | 106.480504 | AA | GG | AG | AG | AA | AA |
| rs13478444 | 107.112543 | AA | GG | AG | AG | AA | AA |
| rs4225396 | 108.071336 | GG | AA | AG | AG | AG | AG |
| rs13478450 | 108.976897 | CC | TT | CT | CT | CT | CT |
| rs13478451 | 109.083781 | GG | AA | AG | AG | AG | AG |
| rs3665124 | 110.024336 | AA | -- | AG | -- | AG | AG |
| rs3708939 | 111.010799 | CC | TT | CT | CT | CT | CT |
| rs32071323 | 111.778516 | AA | GG | AG | AG | AG | AG |
| rs32103915 | 112.501418 | AA | TT | AT | AT | AT | AT |
| rs48981817 | 114.608657 | GG | AA | AG | AG | AG | AG |
| rs29534493 | 115.004548 | TT | CC | CT | CT | CT | CT |
| rs33727786 | 116.673508 | AA | TT | AT | AT | AT | AT |
| rs3023051 | 117.668111 | GG | AA | AG | AG | AG | AG |
| rs13478483 | 118.405516 | TT | CC | CT | CT | CT | TT |

### Supplemental figure 2

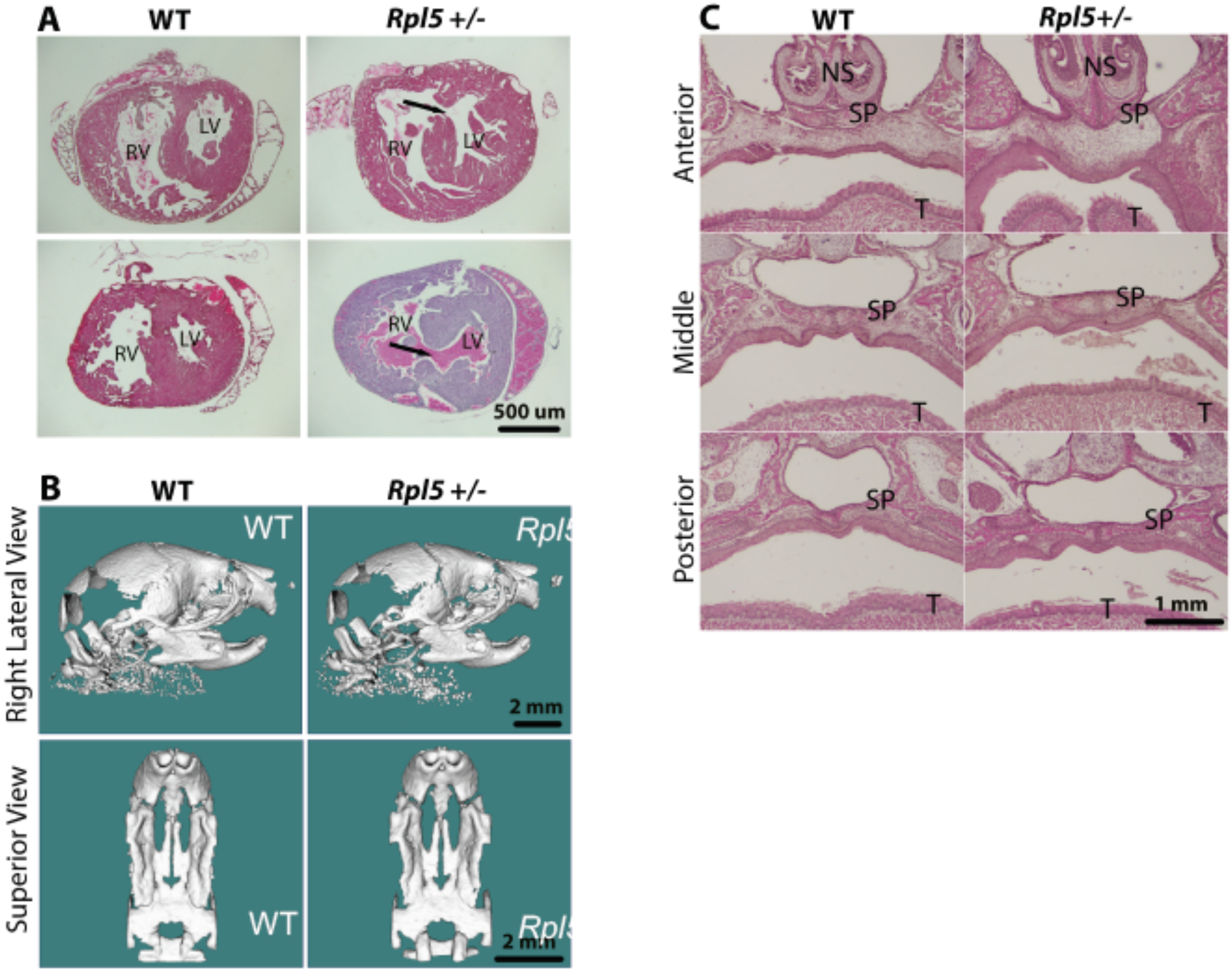


### Supplemental figure 3

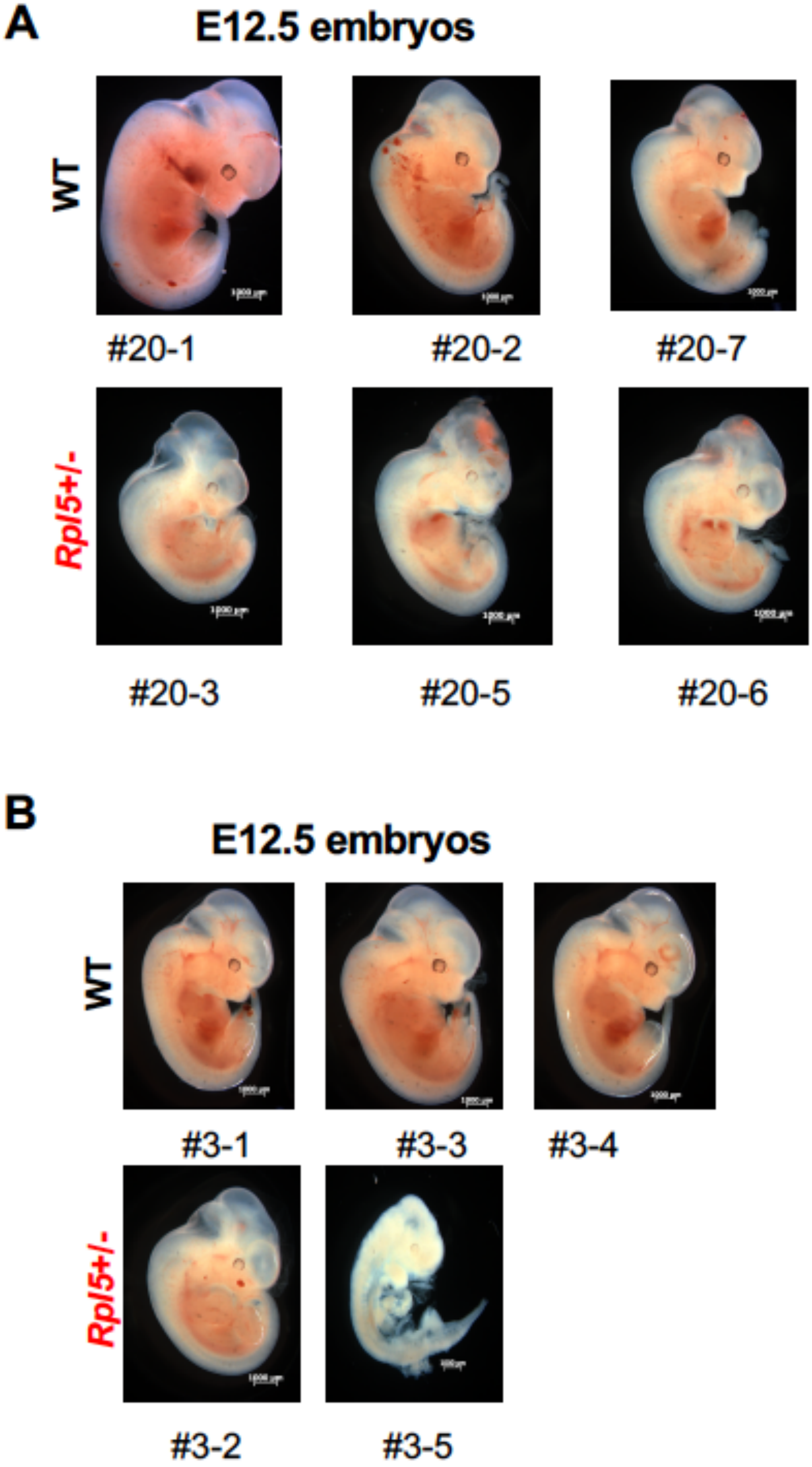


Supplemental figure 4

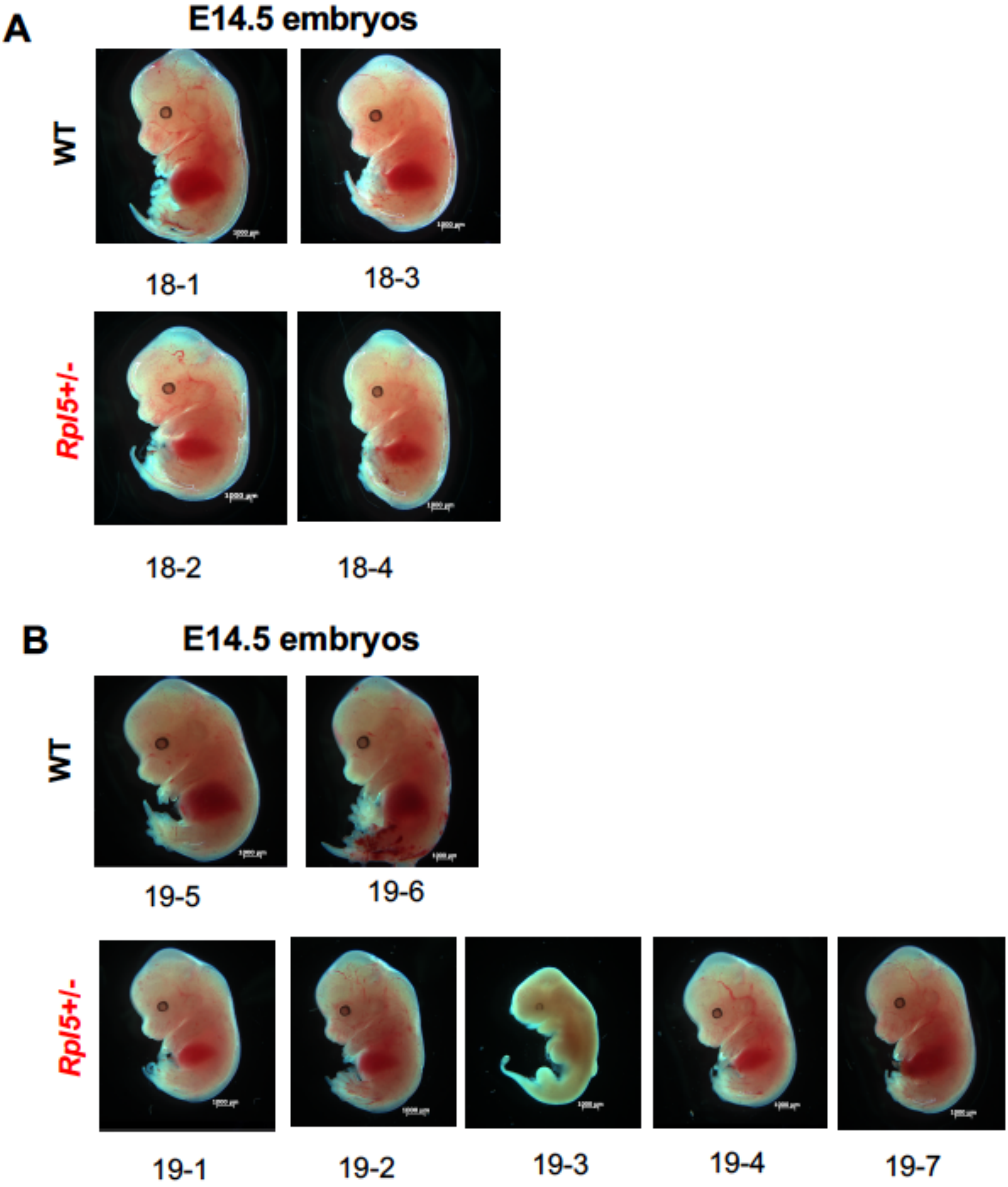


**Supplemental figure 5**

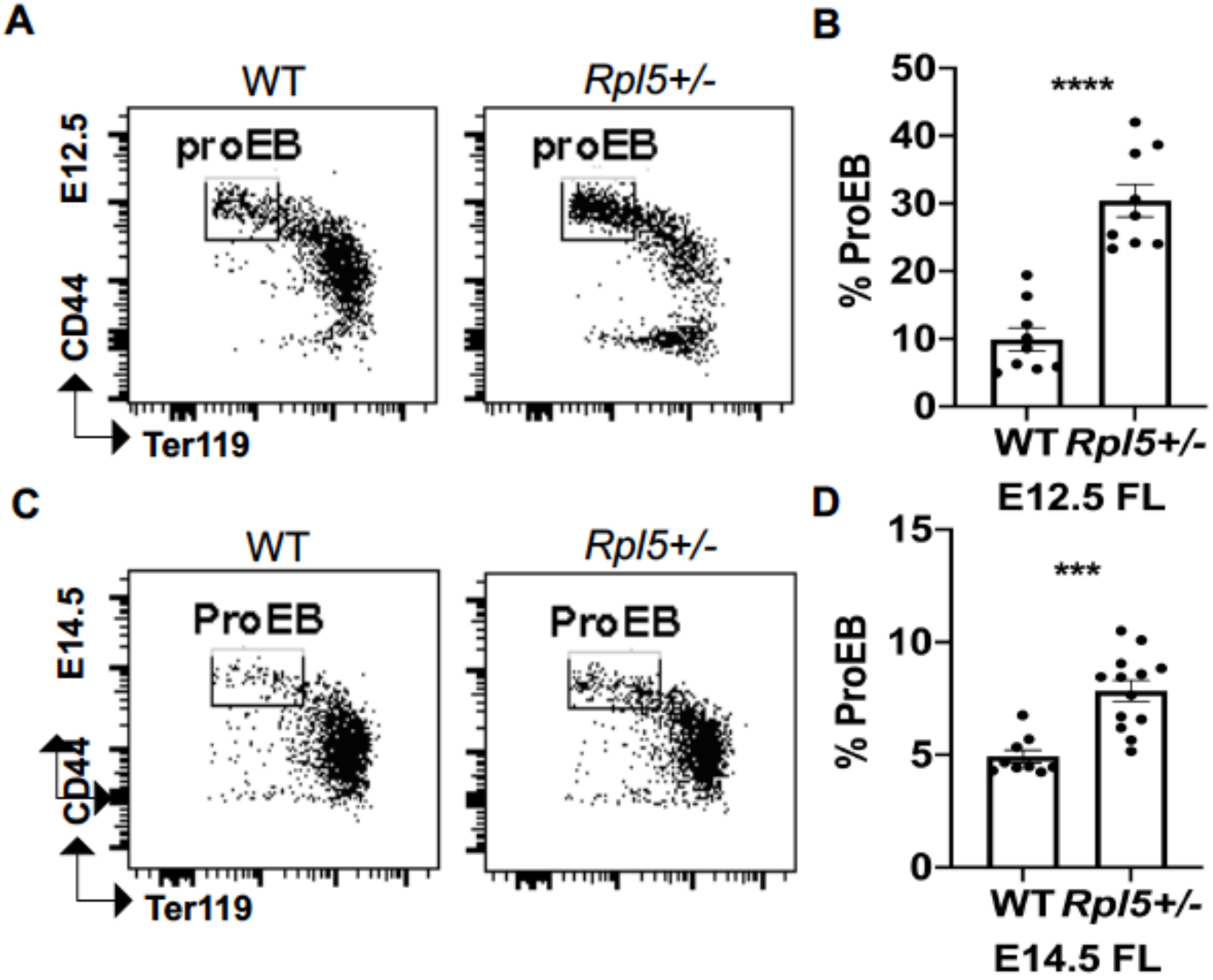


Supplemental figure 6

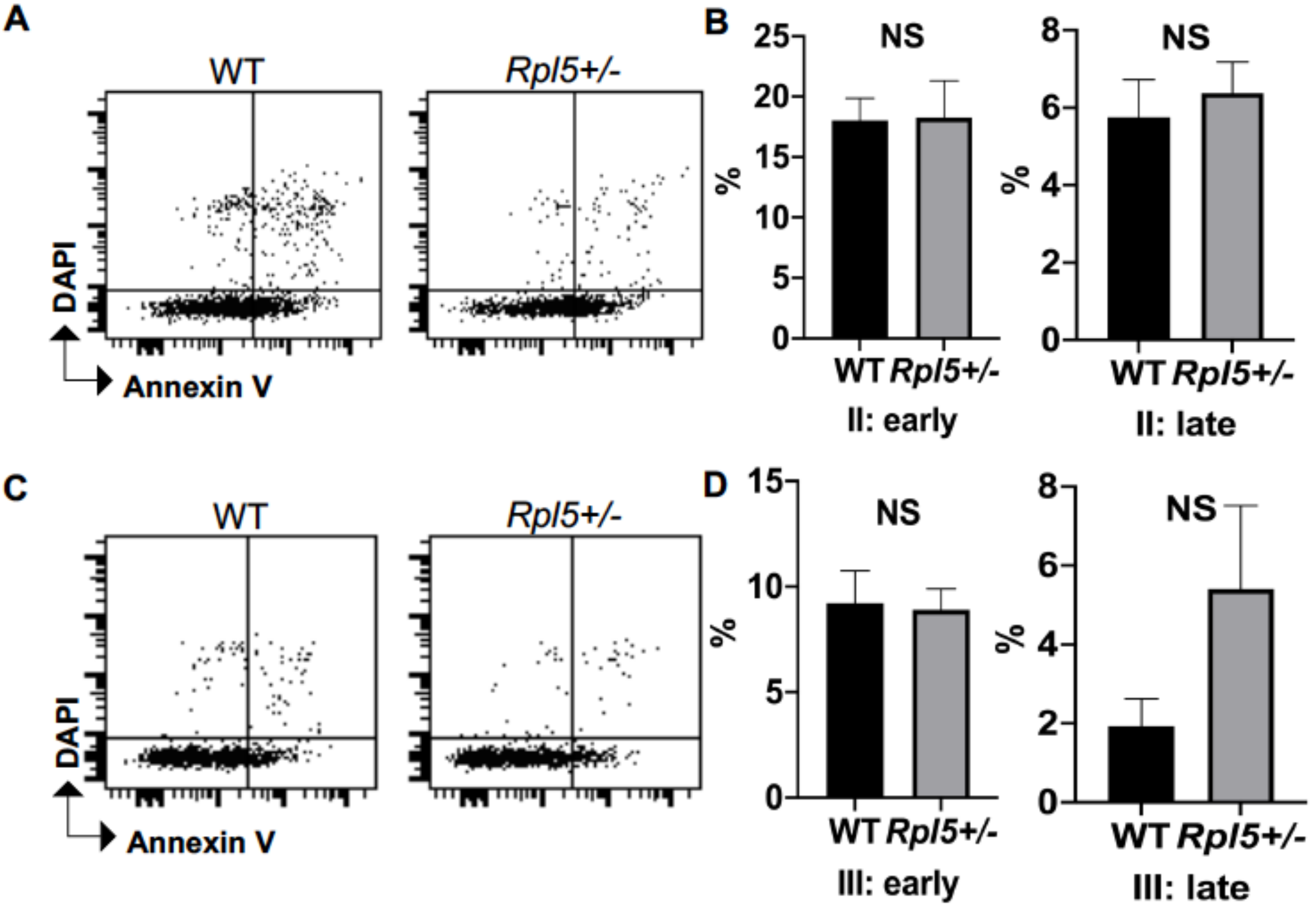


**Supplemental figure 7**

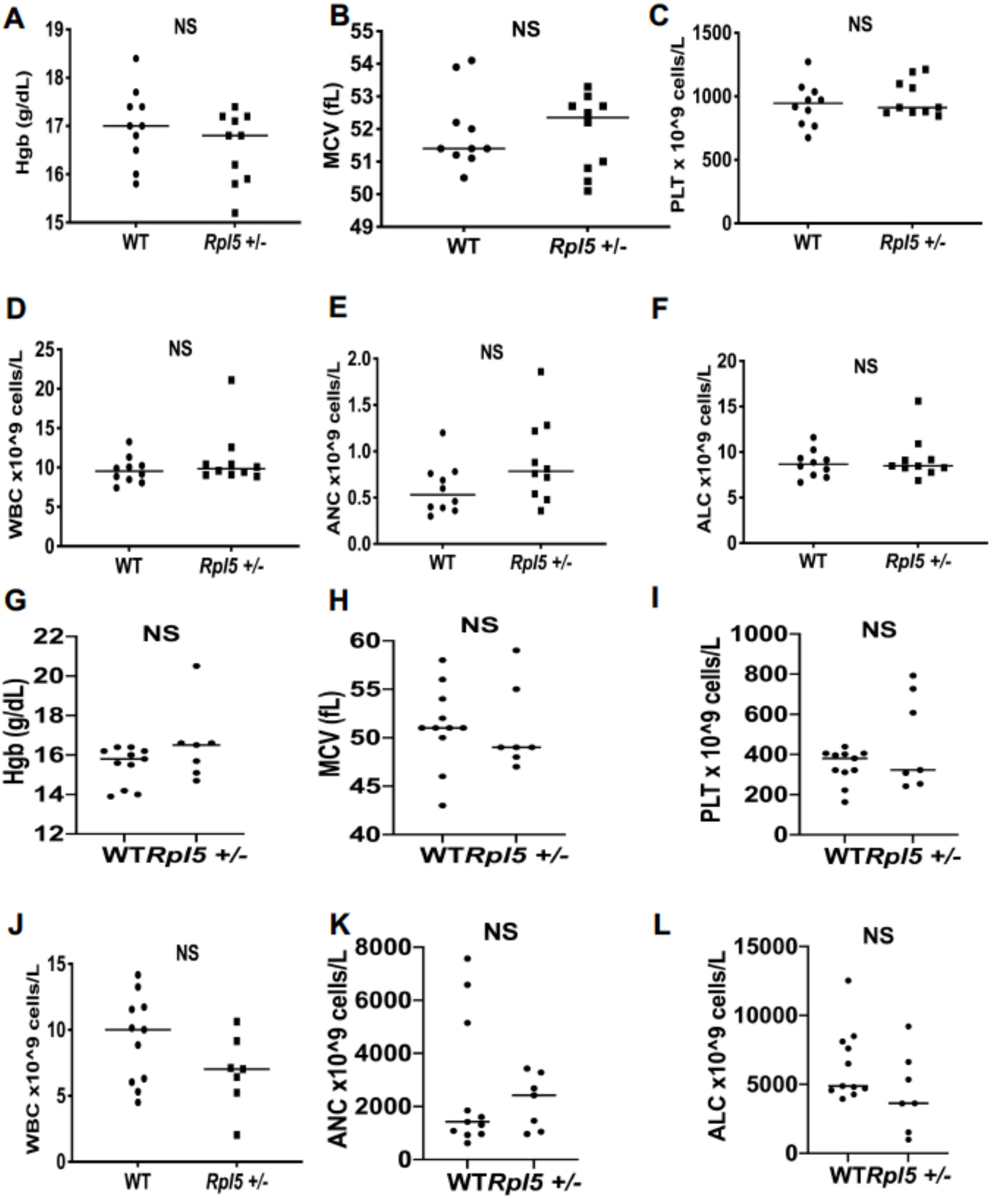


**Supplemental figure 8**

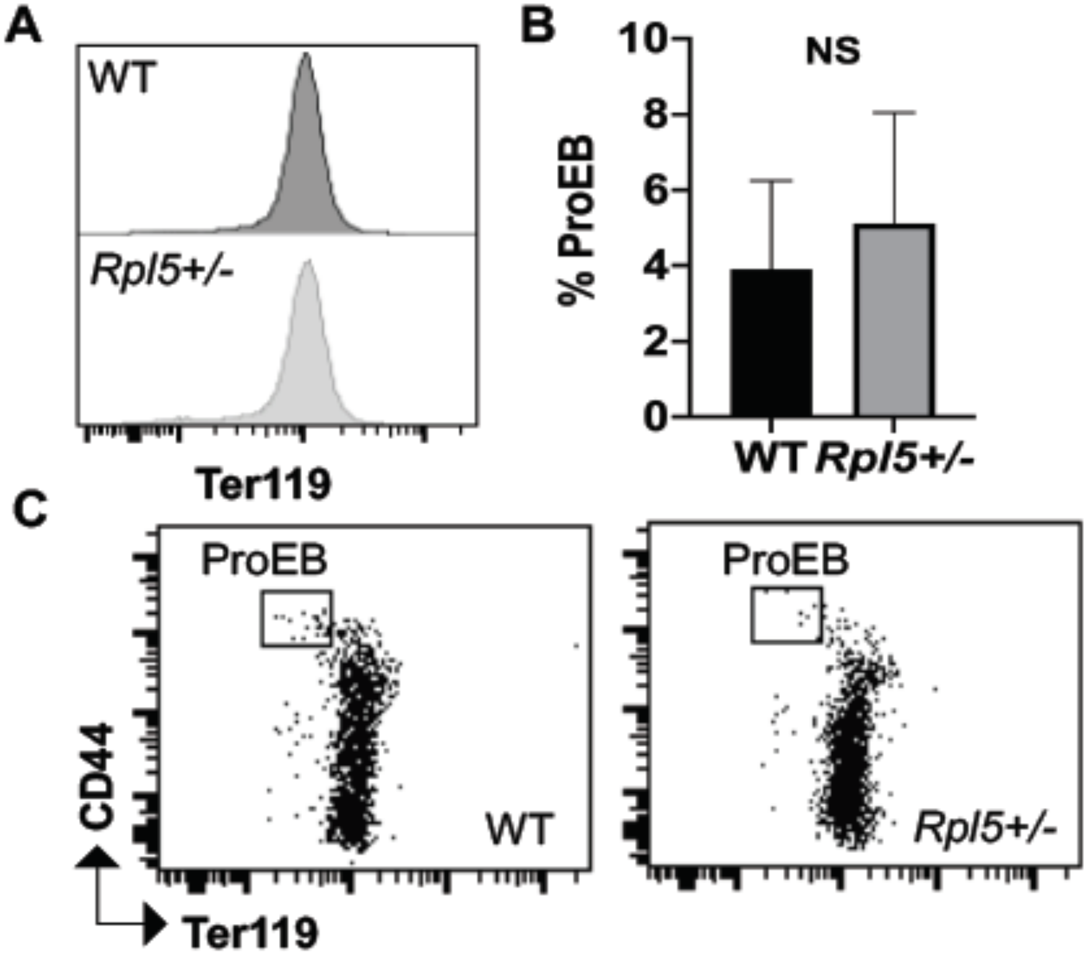


**Supplemental figure 9**

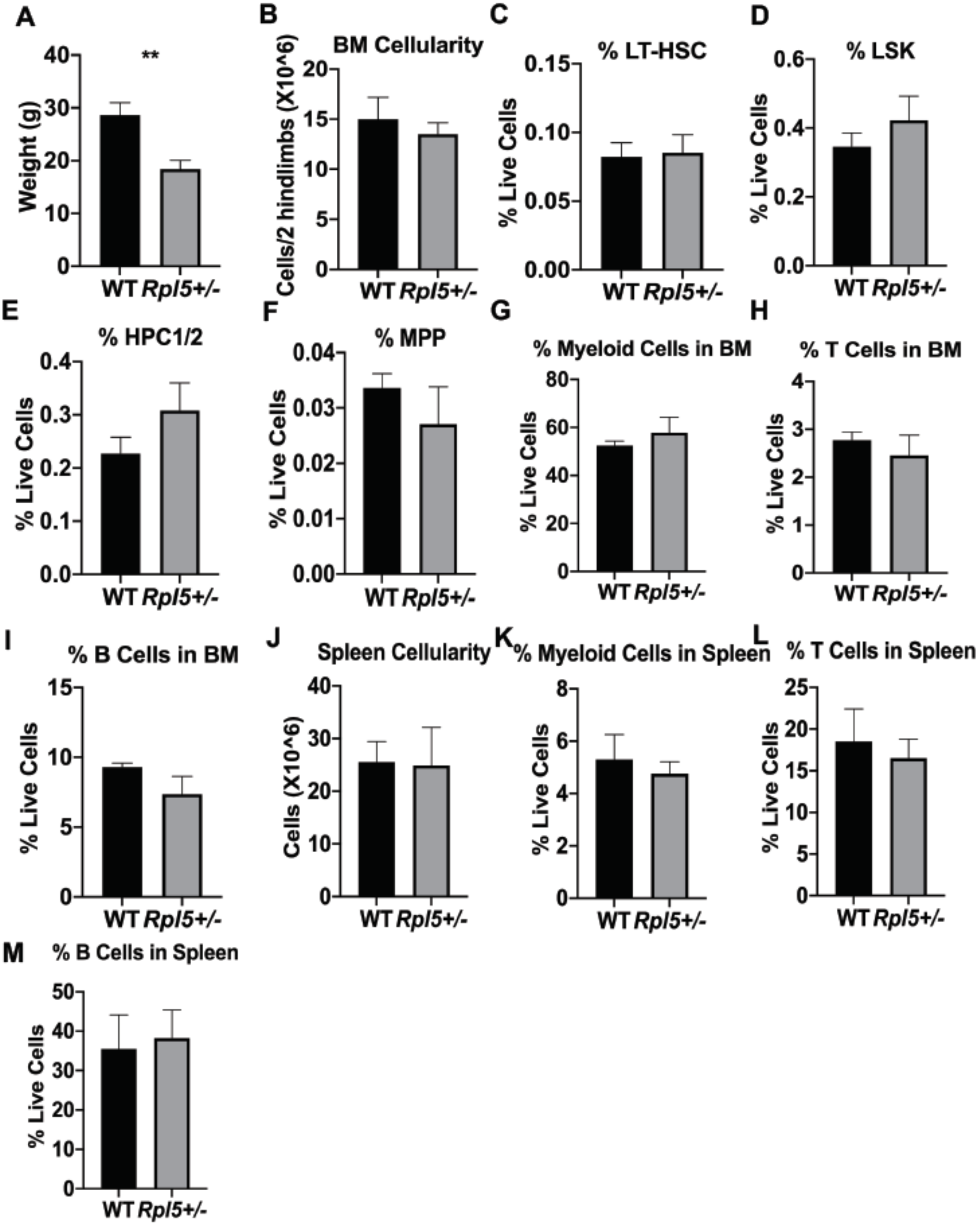


**Supplemental figure 10**
