## Supplemental Information for "A new murine *Rpl5* (*uL18*) mutation provides a unique model of variably penetrant Diamond Blackfan Anemia"

**Supplemental figure 1: Skeletal analysis of neonatal pups.** (A) *Skax23^m1Jus^* pups were smaller in size-smaller mutant pup is shown by white arrow next to 5 wildtype pups of normal size. (B) Whole-mount staining in newborn mice using Alcian blue for cartilage and Alizarin red for bone of WT (left) and mutant mice (right). (C) Compared to bones of WT (wild-type) mice, the length of humeri of *Skax23^m1Jus^* mutant mice was decreased by 19% ± 10.00 (*p* = 0.0927, not statistically significant). Similarly, the length of femurs of *Skax23^m1Jus^* mutant mice was reduced by 16% ± 5.188 (*p* = 0.0169), while the length of their tibia was shorter by 9% ± 5.276 (not statistically significant, *p*= 0.1301). Skulls of mutant mice were smaller anterior-posteriorly than in WT mice (20% ± 11.16, although not statistically significant, p= 0.1207). No significant difference in vertebrae was observed between *Skax23^m1Jus^* mutant mice and WT controls.

**Supplemental figure 2: Identification of the gene mutated in *Skax23^m1Jus^* mutant mice**. (A) Whole genome linkage scans identified a 41 Mb haplotype on chromosome 5 that was heterozygous in B6 in all 21 mice analyzed. The graph shows the percentage of heterozygous mice for the C57BL/6J allele for each SNP (n =149) for each chromosome. B. The table shows the genotypes of these same 21 mice at 13 SNPs selected from the candidate region on chromosome 5. (C) Fine mapping delineated the candidate region on chromosome 5 to an 11 Mb interval flanked by rs13478444 and rs13478483. Heterozygous genotypes at the C57BL/6J allele in panels B and C are shaded.

**Supplemental figure 3: *Rpl5*+/- mice have ventricular septal defect but not craniofacial abnormality.** Mice were euthanized and cardiac and craniofacial sections were generated for histology. (A) No newborn WT mice exhibited cardiac malformations (n=3), while 5/6 mutant mice displayed VSD and 1 mouse also had double outlet right ventricle (DORV; not shown). Analysis of 9 adult mice (WT n=2 and mutant n=7) showed no evidence of VSD by histology. Analysis of the newborn craniofacial structures by (B) micro-CT (WT n=1 and *Rpl5*+/- n=2) or (C) histology (WT n=1 and *Rpl5*+/- n=4) showed no evidence of cleft palate or other craniofacial abnormality.

**Supplemental figure 4: E12.5 embryos**. (A, B) Whole embryo images at E12.5 from 2 litters to demonstrate embryo size and fetal liver color.

**Supplemental figure 5: E14.5 embryos.** (A, B) Whole embryo images at E14.5 from 2 litters to demonstrate embryo size and fetal liver color.

**Supplemental figure 6: Analysis of terminal erythroid differentiation shows a severe defect in E12.5 fetal liver cells.** Analysis of differentiation in E12.5 FL by CD44 vs. Ter119 flow cytometry staining showed a significant block in terminal red cell differentiation with a significant 3-fold increase in the proerythroblast (proEB) population in *Rpl5+/-* animals (A, B). This differentiation block has diminished by E14.5 with a 1.5-fold increase in this proerythroblast block in *Rpl5+/-* mice (C, D). E12.5 WT n=9, *Rpl5+/-* n=9 (3 litters); E14.5 WT n=9, *Rpl5+/-* n=12 (4 litters).

**Supplemental figure 7: Apoptosis in E12.5 FL cells.** E12.5 FL cells were stained with CD71 and Ter119 to analyze red cell differentiation, and then populations II (CD71+ Ter119-/low) and III (CD71+ Ter119+) were analyzed for apoptosis using Annexin V and DAPI staining. No significant difference was observed in early apoptosis (Annexin V+ DAPI-) or late apoptosis (Annexin V+ DAPI+) in the (A, B) II and (C, D) III populations in *Rpl5+/-* versus wildtype embryos. WT=6, *Rpl5+/-* n=6 (2 litters).

**Supplemental figure 8: CBCs in adult and aged mice show no abnormality.** CBCs were performed in young mice (7 weeks old, WT n=10, *Rpl5+/-* n=10; 5 female, 5 male in each group). No significant deviations from normal were observed in (A) hemoglobin (B) MCV (C) platelets (D) WBC (E) ANC (F) ALC for young mice. CBCs were performed in aged mice (WT n= 11, age range 5 – 22 months, *Rpl5+/-* n=7, age range 5 – 17 months). No abnormalities were seen in (G) hemoglobin (H) MCV (I) platelets, (J) WBC, (K) absolute neutrophil count (ANC) or (L) absolute lymphocyte count (ALC). The line represents the median value for each group.

**Supplemental figure 9: Analysis of BM erythroid differentiation profiles in adult animals.** Whole bone marrow was analyzed by flow cytometry for erythroid cell markers (Ter119, CD44) after exclusion of nonerythroid cells (CD45+, Gr-1+, CD11b+, CD3+ and dead cells). (A) No significant difference in Ter119 mean fluorescent intensity or (B, C) in the % of the proerythroblast population (Ter119^low^CD44^hi^) was found between the BM of *Rpl5+/-* and WT adult mutant animals. WT n=3, *Rpl5+/-* n= 3, age 5 months.

**Supplemental figure 10: Analysis of HSPC populations in the BM and spleen of adult animals**. (A) Analysis of hematopoietic populations in WT and *Rpl5+/-* mice was performed by flow cytometry. WT n=5, *Rpl5+/-* n=5 (female mice, age 7-8 months). All counts were normalized by weight (A). No significant defects were seen in: (B) BM cellularity, (C) % LT-HSC (CD150+CD48-LSK), (D) % LSK, (E) % HPC1/2 (CD48+LSK), (F) % MPP (CD150-CD48-LSK), (G) BM myeloid % (CD11b+Gr1+), (H) BM T cell % (CD3+), (I) BM B cell % (B220+), (J) spleen cellularity, (K) spleen myeloid %, (L) spleen T cell % or (M) spleen B cell % populations.

**Supplemental methods:**

### **DNA extraction and genotyping**

Genomic DNA was extracted from mouse tail or embryonic yolk sac using the EZ-10 Spin Column Animal DNA Mini-Preps kit (BS628, Bio Basic Inc. Canada, ON). Genotypes were confirmed for all mice included in the study. Genotyping was done by PCR using genomic DNA and the following primers: an exonic primer 5′ CGTGCTGGTGGTGCTACATA 3′ and intronic primer 5′ GGCCTCCACTGACAGACACT 3′. PCR amplification was done using Q5 High-Fidelity DNA Polymerase (NEB, cat# M0491L) as suggested by the manufacturer. PCR conditions consisted of 30 seconds (s) at 98°C followed by 35 cycles with each cycle consisting of 10s at 98°C, 15s at 67°C, 15s at 72°C, followed by one cycle of 2 min at 72°C. The PCR products were visualized on 1.2% agarose gels. Direct dye terminator sequencing of PCR products was done on an Applied Biosystem's 3730xl DNA Analyzer technology.

**Genotyping by allele-specific PCR**
We established an allele-specific PCR using 3 primers, which were designed using Primer1 software (<http://primer1.soton.ac.uk/primer1.htm>), in standard PCR reactions to specifically identify the *Rpl5* mutant “C” allele. Primers 2FO (forward outer) and 2RO (reverse outer) (5’ CAT CGC CCA TCC TCC ATG CTC GTT GAG 3’ and 5’ CGG CAG CCA TTA GGC CAT GCT GGA AC 3’, respectively) flank a “T” to “C” SNP mutation and amplified a 393 bp amplicon from *Rpl5* wild type and heterozygous genomic DNAs. Primer 2FI (forward inner) sequences (5’ GGG TCT CTA TTC CGC AGG ATG GTG CG**C** 3’) include the “C” allele SNP mismatch (in bold) and another deliberate mismatch at -2 position from the 3’ terminus (underlined). Primers 2FI and 2RO generate a 192 bp amplicon that is specific to the “C” mutant allele. The specificity of these primers was verified using *Rpl5* genomic DNA samples that had been previously confirmed by Sanger sequencing.

**Skeletal studies**

Digital x-rays on adult mice were collected on a Faxitron Ultrafocus instrument. We performed whole-mount Alcian blue and Alizarin red staining of skeleton by euthanizing newborn (day 1-2) mice, removing skin and internal organs and fixing mice in 95% ethanol for 6 days. Samples were then stained overnight in 0.015% Alcian blue solution containing 80% ethanol and 20% acetic acid. Samples were then put back into fresh 95% ethanol for at least 3 hours before transferring into 2% KOH to remove remaining soft tissues. Skeletons were then stained with 0.005% alizarin sodium sulphate in 1% KOH for 12-14 h, then cleared in a solution of 1% KOH and 20% glycerol for at least 3 days and imaged using a dissection microscope (Nikon SMZ 745T). At least 4 newborn mice per group were analyzed for skeletal analysis and data presented as mean ± SD. Linear measurements of skulls and long bones were made using ImageJ software from NIH (version 1.52a). Two-tailed unpaired t-test with equal SD using Graphpad Prism 6.0 (Graphpad Software, Inc.) was used to identify the difference between the two genotypes. The difference was considered statistically significant when *p*-value was less than 0.05.

**RT-qPCR**

Spleen cells were isolated from WT and *Rpl5*+/- mice and lymphocytes were activated in culture with 10 µg/ml LPS for 48 hours. Total RNA was isolated using High Pure RNA Isolation kit (Roche). Reverse transcription was performed using iScript CDNA Synthesis kit (Bio-Rad) and quantitative real-time PCR performed using Power Sybr Green PCR Master Mix (Applied Biosystems/Thermo Fisher). Intron-spanning primers were designed using Primer3web version 4.1.0 (<http://bioinfo.ut.ee/primer3/)>: Rpl5 forward GGCGAGAGGGTAAAACTGAC, Rpl5 reverse ATCCCCTTCTATACGGGCAT, beta actin forward GAGGCATACAGGGACAGCAC, beta actin reverse CTAAGGCCAACCGTGAAAAG. Relative mRNA expression was calculated using the ΔΔCt method and normalized to wildtype cells.

**Blood analysis and flow cytometry**

After euthanasia, blood was drawn by cardiac puncture in adult animals and placed into EDTA tubes. Blood counts were performed using Advia^®^ 120 (Siemans) or VetScan^®^ HM5 Hematology Analyzer (Abaxis). For newborn animals (P1-3), euthanasia was performed by decapitation and blood collected for blood cell counts. Blood samples of newborn animals were diluted 1:10 in PBS prior to analysis in order to obtain sufficient volume to measure. For analysis of hematopoietic stem and progenitor cells, we isolated adult bone marrow and livers of E12.5, E14.5 and P1-3 pups. Cells were incubated with CD16/CD32 Fc block then stained with antibodies in supplemental Table S1. Antibodies for HSPC, mature myeloid and lymphoid analysis are listed in supplemental Table S2. Events were quantified on a Fortessa flow cytometer. Data were analyzed using FlowJo software v. 9.7.6 (TreeStar, San Carlos, CA).

**Cardiac and craniofacial studies**

MicroCTs were performed using μCT100 (Scanco Medical, Bassersdorf, Switzerland) to assess craniofacial structures in adult mice. Three dimensional images were analyzed as outlined in using the MicroView image viewer (Parallax Innovations). For cardiac and craniofacial histology, the tissues were processed, sectioned, and stained by hematoxylin and eosin according to standard protocols. Images were photographed on a Leica DMI3000B microscope using an Olympus DP71 camera and DP controller and manager software.
